# Single-cell transcriptional analysis of lamina propria lymphocytes in the jejunum reveals ILC-like cells in pigs

**DOI:** 10.1101/2023.01.01.522424

**Authors:** Junhong Wang, Ming Gao, Mingyang Cheng, Yu Sun, Yiyuan Lu, Xiaoxu Li, Chunwei Shi, Jianzhong Wang, Nan Wang, Wentao Yang, Yanlong Jiang, Haibin Huang, Guilian Yang, Yan Zeng, Chunfeng Wang, Xin Cao

## Abstract

Pigs are the most suitable model to study various therapeutic strategies and drugs for human beings, while knowledge about tissue- and cell-type-specific transcriptomes and heterogeneity is poorly available. Here, we focused on the intestinal immunity of pigs. Through single-cell sequencing (scRNA-seq) and flow cytometry analysis of the types of immune cells in the jejunum of pigs, we found that innate lymphoid cells (ILCs) existed in the lamina propria lymphocytes (LPLs) of the jejunum. Then, through flow sorting of Live/Dead (L/D)^-^Lineage(LIN)^-^CD45^+^ cells and scRNA-seq, we found that ILCs in the porcine jejunum were mainly ILC3s, with a small number of ILC1s, ILC2s, and NK cells. Through a gene expression map, we found that ILCs coexpressed IL-7Rα, ID2, and other genes and differentially expressed RORC, GATA3, and other genes but did not express the CD3 gene. According to their gene expression profiles, ILC3s can be divided into four subgroups, and genes such as CXCL8, CXCL2, IL-22, IL-17, and NCR2 are differentially expressed. To further detect and identify ILC3s, we prepared RORC monoclonal antibodies and verified the classification of ILCs in the porcine jejunum subgroup and the expression of related hallmark genes at the protein level by flow cytometry. For systematically characterizing of ILCs in the porcine intestines, we combined our pig ILCs dataset with publicly available human and mice ILCs data and identified that the humans and pigs ILCs shared more common features than those mice ILCs in gene signatures and cell states. For the first time, our results showed in detail the gene expression of porcine jejunal ILCs, the subtype classification of ILCs, and the markers of various ILCs, which provides a basis for an in-depth exploration of porcine intestinal mucosal immunity.

Graphical abstract

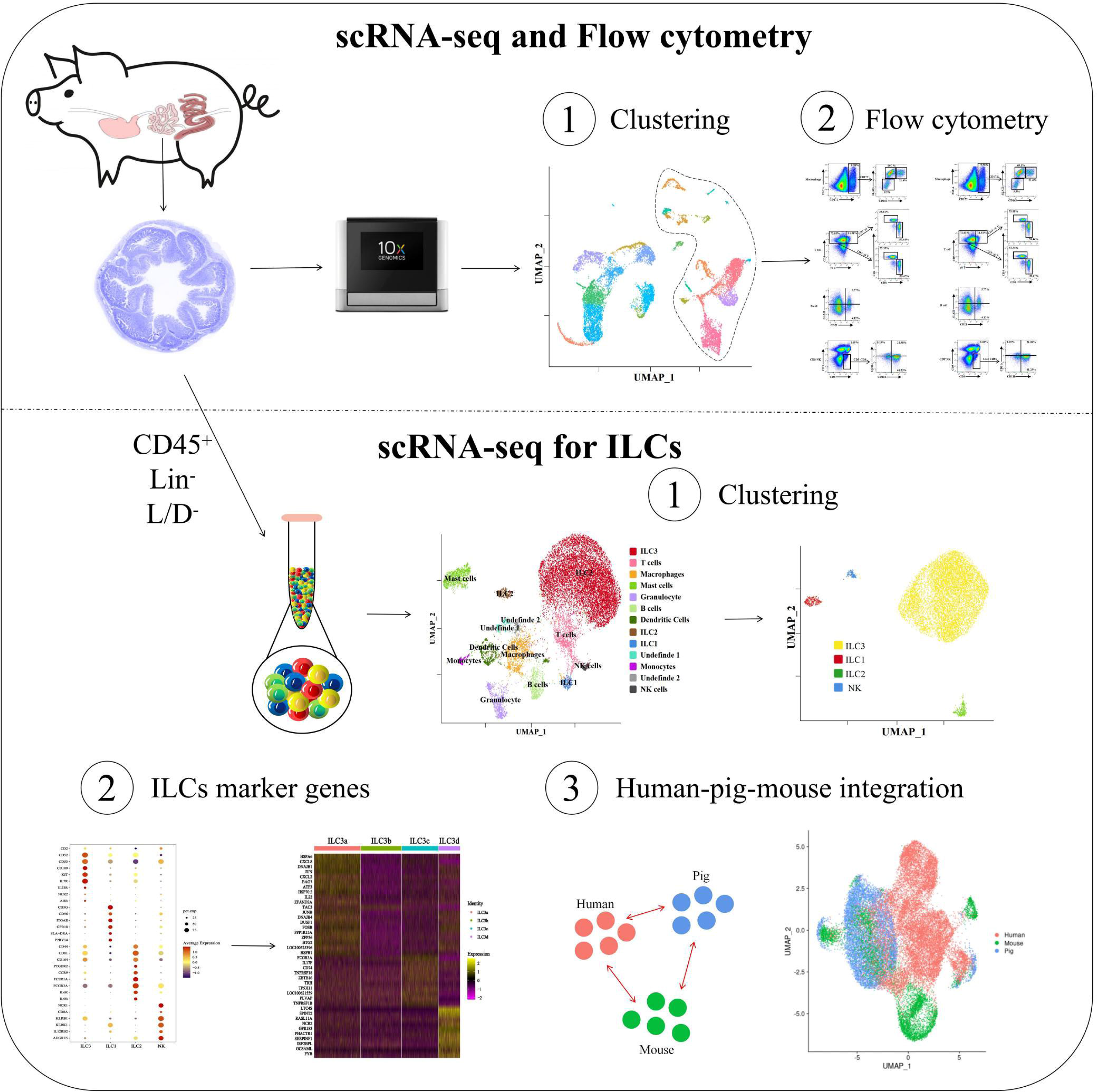

## Introduction

As an ideal animal model that is expected to replace human beings as study subjects, pigs can be used to simulate complex human diseases. For example, pigs can be used as a computational model for predicting human pharmacokinetics^1^, as an animal model for tissue expansion experiments^2^, and as an animal model for burn research^3^. In addition, because they are highly similar to human beings in organ size, structure, anatomy, genetics, and physiological functions, they can be used as ideal donor sources for organ transplantation^4^. Regarding the immune system, the immune cell subsets expressed by pigs are similar to those expressed by humans in phenotype, and the two species have high nucleotide and protein sequence homology in immune molecules^5^. Therefore, pigs have great research value and potential, but basic immunology research related to pigs is still in the primary stage, and there are few reports on the intestinal immunity of pigs, especially on innate lymphoid cells (ILCs).

In recent years, it has been found that innate lymphoid cells (ILCs) play important functions in immunity and tissue homeostasis^6^. The mutual interaction between the microbiome and immune system coordinates the influence of many microorganisms on the host, and ILCs are the frontline that continuously interacts with symbiotic flora in the intestine^7-8^. ILCs are a new family of innate lymphocytes from bone marrow lymphoprogenitor cells and lack classical rearranged antigen receptors^9^. They migrate to various tissue sites and mucous membranes, participating in antipathogen reactions, inflammation, tissue homeostasis, and immune tolerance^10^. According to their functions, four main subgroups of ILCs are defined: ILC1s, ILC2s, ILC3s, and NK cells^11-12^. In addition to ILCs with effector functions, some ILCs with regulatory functions (ILCregs) have been defined^13^. ILC3s have signature gene expressions such as KIT (CD117), IL-7Rα (CD127), and IL-23R^14^. ILC3s can work with commensal microbes to promote intestinal homeostasis and regulate inflammation and mucosal immunity^15^. ILC1s have the closest relationship with Cytotoxic CD8^+^ T cells, expressing higher levels of genes encoding cytotoxic molecules than other ILC/T types^16^. A previous study found higher levels of Th2/ILC2-type cells in the lung mucosa^17^. ILC2s have functional similarities to Th2 cells in terms of the expression of the master transcription factor GATA3 and the production of functional cytokines and can play important roles in the expulsion of parasites and induction of allergy through secretion of IL-5 and IL-13, including myeloid cell recruitment, eosinophil maturation, smooth muscle contraction, and mucin production by epithelial cells^18^. To date, NK cells are the only identified porcine non-B/non-T-lymphocyte subset and are an important innate immune system cell population. With the ability to attack natural pathogens with infected and malignant body cells and produce immunomodulatory cytokines, NK cells are commonly identified by flow cytometry as CD3ε^-^CD8α^+19^.

Single-cell RNA sequencing (scRNA-seq) has been used to describe the gene expression of immune cells in many tissues and organs of pigs, including peripheral blood, thymus, lung, and embryonic tissue^20-21^. The heterogeneity of endothelial cells, microglia regulation, and transcriptome changes under a single-cell resolution of virus-infected porcine macrophages were also revealed through the cell landscape at the pig single-cell level^22^. Recently, researchers have found suspected ILCs in pigs’ intestinal immune cells^23^. However, there is no report on the gene expression and cell typing of ILCs in the porcine intestine. We extracted total lamina propria lymphocytes (LPLs) from the porcine jejunum, obtained Live/Dead (L/D)^-^Lineage(LIN)^-^CD45^+^ cells (amount: 200,000) by fluorescence-activated cell sorting (FACS), and then performed scRNA-seq. We recorded and presented the ILC population and heterogeneity level in the porcine jejunum, which was not previously described. Collectively, these data are the highest resolution transcriptome maps of ILCs in the intestinal immune landscape of pigs and can be used to further decode the phenotype and function of ILCs in the intestinal tract.

Our data provide a resource better to understand porcine ILCs of LPLs in the jejunum. By performing scRNA-seq on immune cells from the jejunum of piglets, we recorded and detailed the specific typing of porcine ILCs and the characterizing of the ILC3 subgroup. It was found that there was a certain transdifferentiation process of ILCs in the piglet jejunum under complex intestinal conditions. Cluster analysis of ILCs in humans, pigs, and mice showed that the gene expression similarity of human and pig intestinal ILCs was high, but the similarity between pigs and mice was not high, which suggested that swine could be a better substitute for humans as model animals to investigate intestinal mucosal system homeostasis after various challenges.

## Results

### 1. Characterization of immune cells in LPLs of the jejunum in pigs

The immune cells in the porcine jejunum lamina propria are abundant, and the cell types are obvious. We identified a large number of myeloid cells present in the pig lamina jejuna, including dendritic cells (CD172^+^SLAII^+^CD163^-^), macrophages (CD172^+^SLAII^+^CD163^+^), polymorphonuclear neutrophil (CD172^+^SLAII^-^CD163^-^), and monocytes (CD172^+^SLAII^+/-^CD163^+/-^). T cells in the lamina propria include two cell subsets, namely, CD3^+^γδT^-^ and CD3^+^γδT^+^, and we found more CD4^+^ cells in the CD3^+^γδT^-^ subset and more CD8^+^ cells in the CD3^+^γδT^+^ cell subset. We found a certain proportion of B cells expressing SLAII and CD8^+^ NK cells expressing CD11b, and CD11c were present in the pig lamina propria^24^. To obtain a large number of ILCs, the cells obtained by flow sorting were subjected to subsequent single-cell transcriptome sequencing (Figure S1A).

The analysis of single-cell transcriptome data of the pig jejunum shows that there are a large number of nonimmune cells and immune cells in the jejunum, among which immune cells (CD45) are mainly divided into myeloid cells, T cells (CD3), B cells (CD21) and ILCs. However, at this resolution, ILCs are mainly ILC3s (IL-23R), and the classification of ILCs into other subgroups is unclear. These immune cells constitute the porcine jejunum’s immune system (Figure S1B, C) (Table S1).

To better explore the specific grouping information and expression profiles of inherent lymphoid cells in the jejunum of pigs, we removed the cells with abnormally low and high gene counts and high mitochondrial gene expression and then used Seurat to cluster the remaining cells^25^. A total of 15,763 immune cells were obtained. According to the different transcription levels of genes of different cells, they can be preliminarily divided into four types of cells (Figure 1A), among which there are 11,208 ILCs/T cells, whose surface receptors are CD127 and nuclear transcription factors such as GATA3 and RORC. 3,500 myeloid cells express THBS1, TIF1, MS4A2, and other landmark genes of myeloid cells, and the gene expression of myeloid cells is heterogeneous, which is related to the functional polarization of myeloid cells. 717 B cells express B-cell markers such as MZB1, JCHAIN and TNFS17R, and 338 unknown immune cells (Figure 1B, 1C). According to the gene expression clustering UMAP analysis, the cells were divided into 13 cell subsets (Figure 1D) (Table S2), in which the symbolic marker expressed in each subgroup is shown in Figure 1E. With CD172 antibodies in Lin, the single-cell results showed that only some myeloid cells were detected, such as the cell subsets of macrophages, mast cells, granulocytes, dendritic cells, and monocytes. Although we removed most of the non-ILCs from the immune cells, some unknown cells were still present, suggesting that the immune cell subsets in the pig intestinal lamina propria are enriched, possibly due to the influence of the complex intestinal environment.

**Fig. 1.**
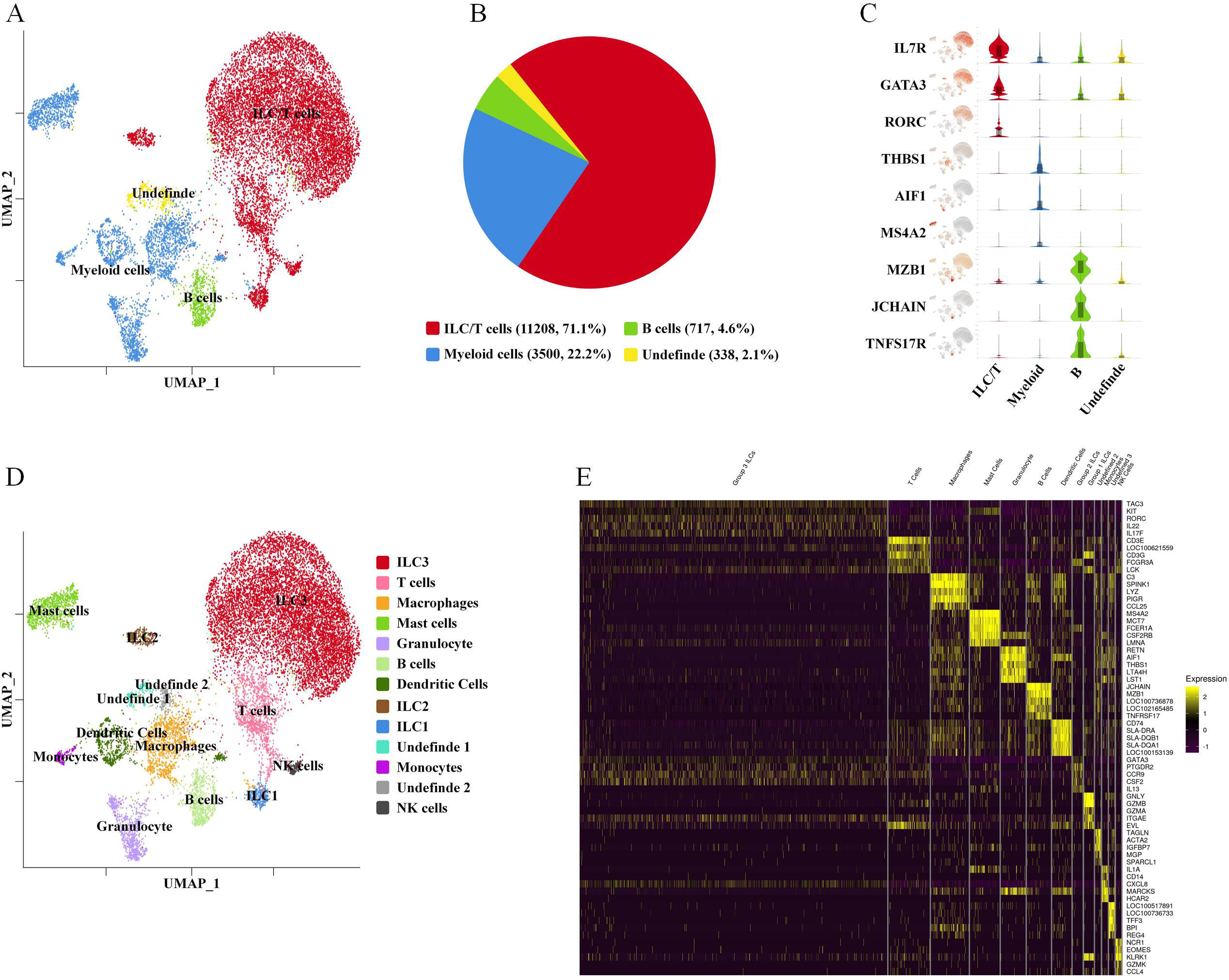
Characterization of immune cells in the Pig jejunum after flow cytometry sorting. A: UMAP plots showing the immune landscape of the 4 subpopulations of the intestine cell(Selection by flow cytometry). Cells are color-coded according to the defined subset(ILC/T cells: red; Myelid cells: blue; B cells: green; Undefined cells: yellow). B: Pie charts showing quantitative differences for these four classes of cell clusters, ILC/T cells: 11208 cells(71.1%); Myelid cells: 3500 cells (22.2%); B cells: 717 cells(4.6%); Undefinde: 338 cells(2.1%). C: Violin plot representation of differentially expressed marker genes for the four clusters, Three signature genes were displayed in each cluster, except for the Undefined cells. D: UMAP plots showing the immune landscape of the 13 clusters of intestine cells. E: Heatmap showing the marker genes with the five highest differentially expressed in each cluster, Abscissa shows different clusters, and ordinate shows differentially expressed gene names.

### 2. ILC3s are the most abundant cell type among Lin^-^CD45^+^ cells in LPLs of the jejunum in pigs

Early sorting has removed many T cells and is not abundant for typing T-cell subpopulations. The regrouping results showed that ILC/T cells could be divided into seven subsets according to their hallmarks, namely, ILC3s, CD8^+^ T cells, CD4^+^ T cells, CD3^+^ T cells, ILC1s, ILC2s, and NK cells (Figure 2A). Studies have shown that specific cytokines and transcription factors produced by Th1, Th2, and Th17 cells are also secreted by ILC1s, ILC2s, and ILC3s^26^, respectively. The results of the surface molecular responses of T cells and ILCs were quite different. T cells expressed CD3E, which could be divided into CD4 T cells and CD8 T cells, while ILCs expressed surface markers such as IL-23R (ILC3), PTGDR2 (ILC2), and NCR1 (NK) (Figure 2B). Both T cells and ILCs express the chemokines CXCL8 (ILC3, CD8 T, and CD3 T cells) and CCL5 (CD8 T, CD3 T, ILC1, and NK cells), and CD4 T cells express more IL-10 (Figure 2C). For transcription factors, T cells and ILCs are similar, as both express ID2 and RORC (ILC3, CD3 T); IKZF3 is mainly expressed by T cells, ILC1s, and NK cells; TBX21 and EOMES are mainly expressed by NK cells; and GATA3, as an important regulator of T-cell development, is expressed in ILC/T cells (Figure 2D).

**Fig. 2.**
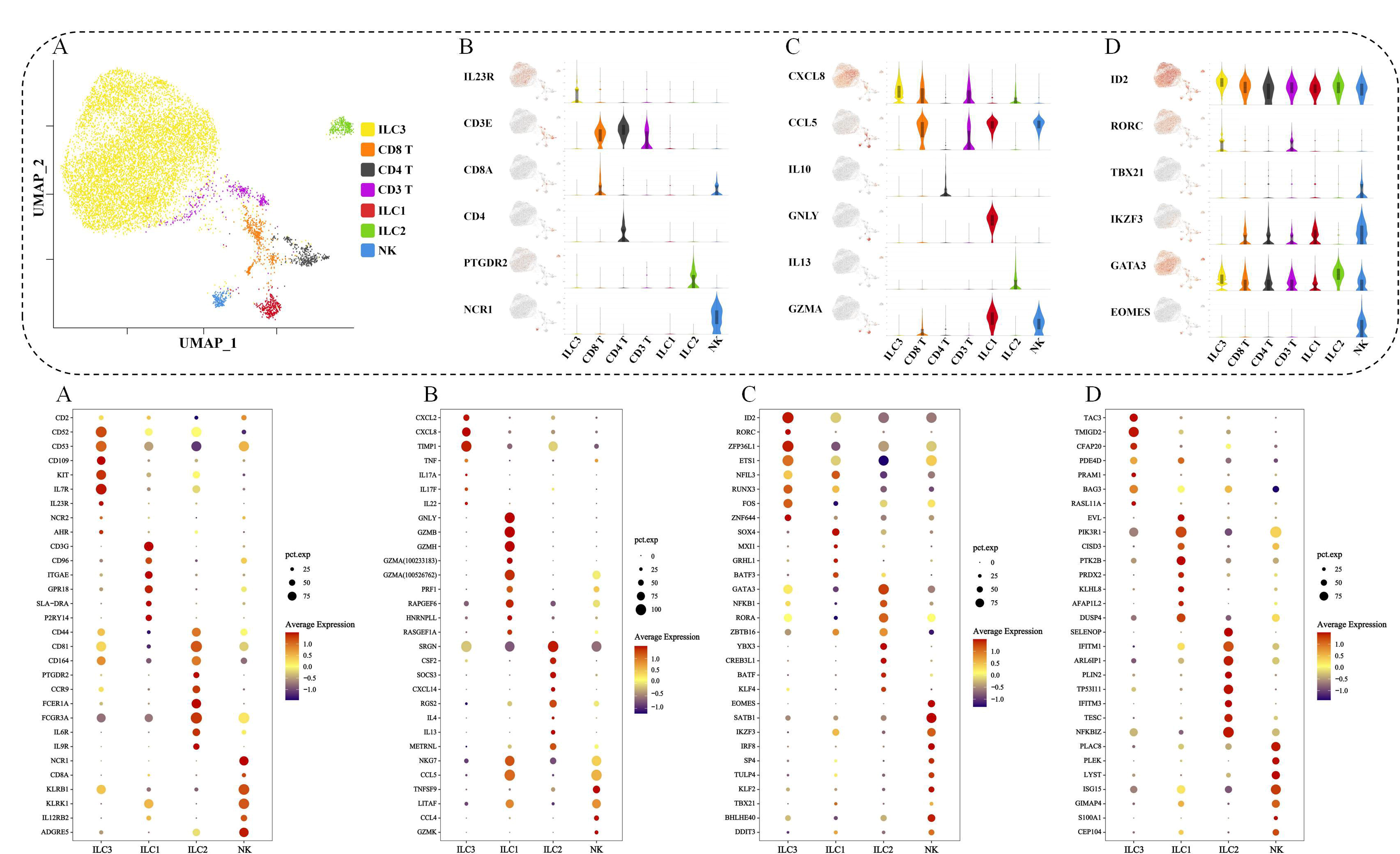
Detailed typing of ILCs in pig jejunum. A: UMAP plots showing the immune landscape of the 7 clusters of intestine cells (ILC/T cell regrouping). Violin statistics of the marker genes for ILC and T cell differential expression. B: Receptor protein genes. C: Secreted protein genes. D: Transcription factor genes. Detailed statistics of the expressed genes in the four subsets of the ILC. Dot plot showing Z-scored mean expression of selected marker genes in ILC clusters from, the size of the dot represents the percentage of the gene in a subset of cells, The colors of the dots represent the average expression of the gene in a subset of cells. E: receptor protein genes. F: Secreted protein genes. G: Transcription factor genes. H: other protein-encoding genes.

In the barrier tissues of mammals, ILCs account for approximately 0.5-5% of lymphocytes, so they are a rare cell group^27^. Our scRNA-seq results showed that there were ILCs in the intestinal lamina propria of pigs, which accounted for a certain proportion (Figure S2A). Cluster analysis of ILCs showed that the correlation between ILC3s and ILC2s was higher, while that between NK cells and ILC1s was higher (Figure S2B). For the GSVA analysis of ILCs (Figure S2C) (Table S3), subsets of ILCs were highly enriched in pathways such as LINOLEIC_ACID_METABOLISM, RENIN_ANGIOTENSIN_SYSTEM, and MATURITY_ONSET_DIABETES_OF_THE_YOUNG, suggesting that ILCs play a crucial role in the development and are involved in the regulation of morphogenesis in various tissues and organs. However, the enrichment degree in PENTOSE_PHOSPHATE_PATHWAY, ACID_METABOLISM, and other pathways was relatively low.

We performed a detailed statistical analysis on the surface receptors, effector factors, transcription factors, and other highly expressed genes of ILCs (Table S4). We found that, except for ILC3s, other ILCs accounted for a relatively low proportion of the total ILCs.

#### ILC3s

We found that porcine ILC3s expressed many immune characteristic genes of this cell type. For the surface receptors, ILC3s expressed iconic surface receptors such as CD2, KIT, IL-7Rα, IL-23R, NCR, and AHR (Figure 2E). IL-7Rα plays an important role in developing and differentiating ILC3s^28^, and NCR2+ ILC3s uniquely maintain intestinal homeostasis^29^. ILC3s highly expressed the powerful chemokines CXCL2 and CXCL8 of neutrophils (Figure 2F), which revealed their regulatory effect on granulocyte-affecting migration and survival^30^. Like TH17 cells, ILC3s express the TNF, IL-22, IL-17A, and IL-17F genes, which is why ILC3s play an important role in the innate immune response, especially in the intestinal mucosa^31^. ILC3s represent a series of transcription factors (Figure 2G), among which ID2 is an important transcription factor that regulates many cellular processes^32^; RORC (RORγt) has been used as a symbolic transcription factor of ILC3s^33^; and Runx3 is necessary for the normal development of ILC1s and ILC3s. ETS1 is involved in stem cell development, cell senescence, and death^34^, and FOS regulates cell proliferation^35^, differentiation, and transdifferentiation. We also found that ILC3s in the porcine intestine highly expressed the gonadotropic development-related gene TAC3 (Figure 2H)^36^, and the TMIGD2 gene was involved in the positive regulation of T-cell activation^37^. The transcribed genes of ILC3s were enriched in pathways such as HEMATOPOIETIC_CELL_LINEAGE, NITROGEN_METABOLISM, CYTOKINE_CYTOKINE_RECEPTOR_INTERACTION, and ARACHIDONIC_ACID_METABOLISM. ILC3s are involved in innate and adaptive inflammatory host defense, cell growth, differentiation, cell death, angiogenesis, and developmental and repair processes to restore homeostasis and regulate nitrogen metabolism, arachidonic acid metabolism, and other processes (Figure S2D).

#### ILC1s

ILC1s still express other genes related to the CD3 complex, such as CD3G and CD247 (encoding CD3z) (Figure 2E). The antigen presentation-related genes CD96 and SLA-DRA and the gene ITGAE (integrin subunit αE), which mediates the adhesion of T lymphocytes to epithelial cells, were also expressed. The expression of genes encoding cytotoxic molecules, including GZMA, GZMB, GZMH, and GNLY, was significantly increased (Figure 2F). Additionally, the gene PRF1 encoding the perforin protein was found. The main regulator gene of activation-induced alternative splicing in T cells is HNRNPLL^38^. The transcription factors identified for ILC1s include SOX4 and the transcription inhibitor MXI1 (Figure 2G). It was also found that ILC1s were highly responsive to the transcription factor GRHL1 and the transcription inhibitor BATF3. The EVL gene had high expression in T cells and ILC1s (Figure 2H). It has been reported that the PRDX2 gene may contribute to the antiviral activity of CD8+ T cells^39^. The ILC1-enriched pathways are all related to autoimmune diseases, inducing cytotoxic T lymphocyte (CTL) and natural killer (NK) cell responses (Figure S2D).

#### ILC2s

Our results showed that ILC2s expressed TH2/ILC2 surface marker genes, including CD44, CD81, and CD164 (Figure 2E), which regulate cell development, activation, growth, movement, adhesion, and migration. ILC2s also expressed PTGDR2; the chemokine receptor genes CCR9; interleukin-6 receptor (IL-6R) and interleukin-9 receptor (IL-9R); the receptor gene Fcgr3a encoding the fc part of immunoglobulin g; and the IgE receptor encoding gene FCER1A. ILC2s express TH2 cytokines such as IL-4, IL-5, and IL-13 and express CSF2 (Figure 2F), which controls the production, differentiation, and function of granulocytes and macrophages. ILC2s also express the cytokine signaling inhibitory gene SOCS3 and the chemokine gene CXCL14, which are involved in the homeostasis of macrophages. ILC2s significantly expressed GATA3, NFKB1, RORA, ZBTB16, YBX3, CREB3L1, BATF, KLF4, and other transcription factors (Figure 2G). ILC2s expressed the negative regulatory gene TP53I11, which is involved in cell population proliferation (Figure 2H)^40^; the TESC gene, which regulates molecular metabolic processes and bone marrow cell differentiation^41^; the IFN-induced antiviral protein family genes IFITM1 and IFITM3; and NFKBIZ, which encodes the nuclear I kappa B protein. The gene sets enriched in ILC2s are related to intestinal immunity. A significant feature of intestinal immunity is that it can produce many noninflammatory immunoglobulin A (IgA) antibodies, which can serve as the first line of defense against microorganisms (Figure S2D).

#### NK cells

NK cells of pigs expressed CD8a, CD11b, and NCR1 (Figure 2E). These cells express several genes (killer cell lectin-like receptor) of superfamily C, KLRB1, KLRFI, and KLRK1; the IL-12 receptor gene IL-12RB2; and ADGRE5, which mediates cell-cell interactions. In terms of cytokines, we found that NK cells are similar to CD8 T cells/ILC1s (Figure 2F), both of which express NKG7 and the chemokine genes CCL4 and CCL5, which are involved in immune regulation and inflammatory processes^42^. In terms of transcription factors, NK cells expressed the transcription factors EOMES and TBX21 (coexpressed by ILC1s and NK cells) (Figure 2G)^43^; the transcription factors IKZF3 and IRF8^44-45^, which regulate lymphocyte development; KLF2, which is associated with early animal development^46^; a gene that controls cell differentiation, namely, BHLHE40; and the apoptosis-related gene DDIT3^47^. On the other hand, NK cells also express PLAC8, PLEK, LYST, ISG15, GIMAP4, S100A1, CEP104 and other genes (Figure 2H). NK cells were shown to be enriched in pathways such as KEGG_SYSTEMIC_LUPUS_ERYTHEMATOSUS and KEGG_GLYCOSPHINGOLIPID_BIOSYNTHESIS_LACTO_AND_NEOLACTO_SERIES (Figure S2D).

According to the statistical results of interacting genes among ILC subsets, we found that there were 32 ligand binding relationships among ILC subsets (Figure S3A, B), the most significant of which was the obvious interaction between NK cell subsets and other ILCs, which also proved the important relationship between NK cells and antitumor, antiviral infection and immunomodulation (Figure S3C). The obvious ligand CD74-MIF plays an important role in cell-mediated immunity, immune regulation, and inflammation and can produce anti-inflammatory effects by inhibiting glucocorticoids and regulating macrophage function in host defense. EPHB6-EFNB1 may play a role in cell adhesion and the development or maintenance of the nervous system^48^. The binding of TNF- and its receptor is involved in the response of cells to stimuli such as cytokines and stress and plays a key role in regulating the immune response to infection^49^. C5AR1-RPS19 participates in the biochemical pathway of the innate and adaptive immune response-complement system^50^. CD44-HBEGF is involved in various cellular functions, including activation, recycling, and homing of T lymphocytes; hematopoiesis; inflammation; and response to bacterial infection^51^.

### 3. Cluster analysis of ILC3s

To classify and analyze the subsets of ILC3s in more detail, we found that the ILC3 group can be divided into four subgroups (Figure 3A) according to their gene expression differences. The correlation analysis results show that ILC3b and ILC3c have a high correlation, while ILC3d has a low correlation with other ILC subgroups (Figure 3B). Gene expression profiles of the ILC3 gene list show the four subsets (Figure 3C) (Table S5), and significant differences in the transcription levels among the ILC3 subgroups were found (Figure 3D). The high expression of the heat shock protein genes HSPA6 and DNAJB1 (Figure 3E) in the ILC3a subset suggests that ILC3a cells may play an important role in maintaining the normal function of intestinal cells, thus regulating the balance between the survival and death of cells. The expression of inflammatory cytokine genes such as CXCL2, CXCL8, and IL-22 suggests that this cell subset plays an important role in the inflammatory response.

**Fig. 3.**
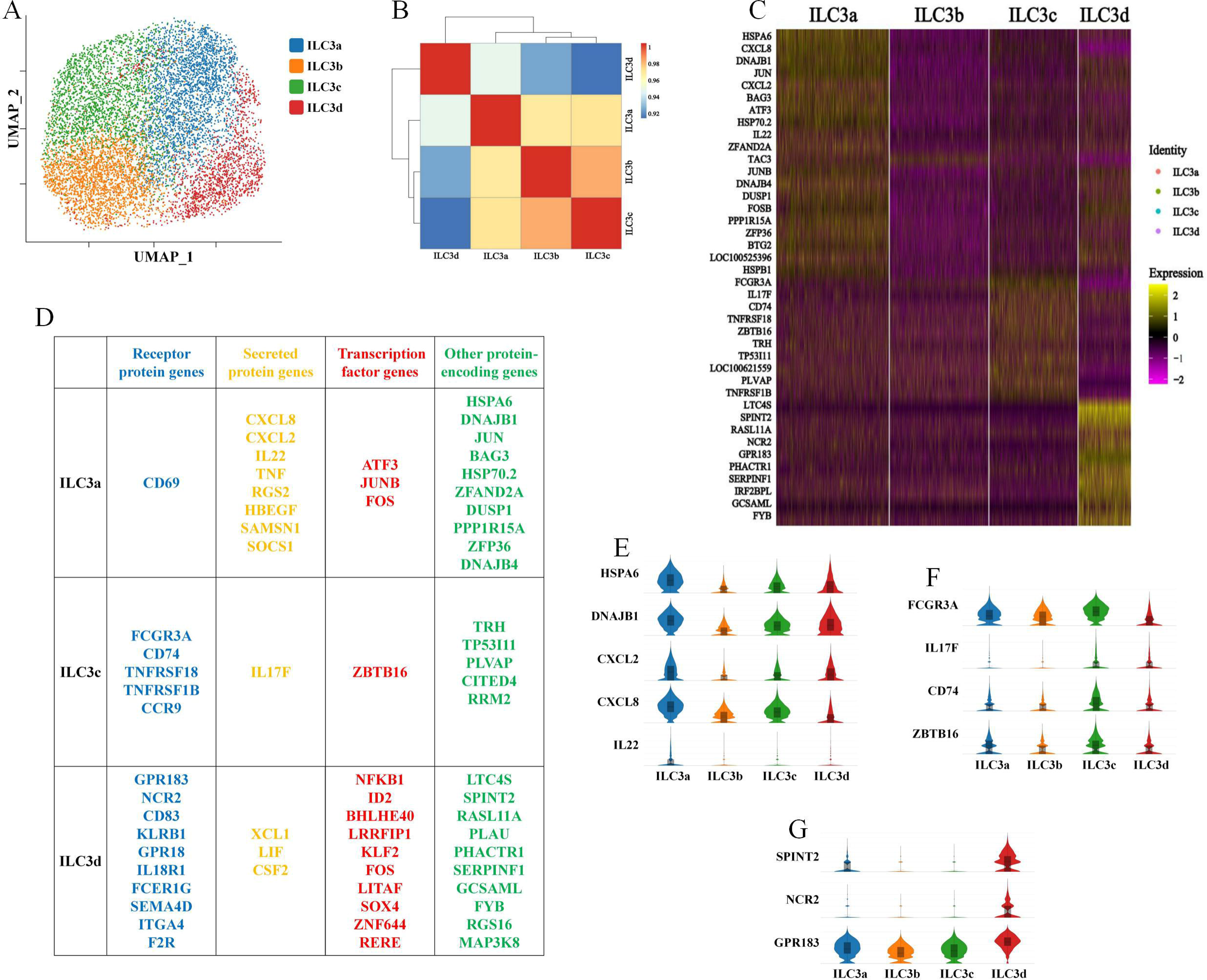
Detailed typing research of ILC3 in pig jejunum. A: UMAP plots showing the immune landscape of the ILC3 clusters. Cells are color-coded according to the defined subset(ILC3a cells: blue; ILC3b: orange; ILC3c cells: green; ILC3d cells: red). B: Spearman Correlation Analysis between subsets of ILC3, Colors represent the degree of correlation between the ILC3 clusters, and Tree branches represent correlation cluster grading. C: Heatmap is the differential genes’ average expression of ILC3 clusters. The top 10 differential genes were taken from the ILC3a, ILC3b, ILC3c, and ILC3d. Among these, only one differential gene in ILC3 b cells was upregulated, while the remaining differential genes were downregulated. D: Top ten expressed genes encoding receptor protein(blue), secreted protein(orange), transcription factor (red), and other protein-encoding(green) significantly differentiating among the groups of ILCs. Gene symbols and annotations were retrieved from public databases. Genes are ranked by *P*-value. E: Violin plots of the ILC3a highly expressed genes. F: Results of the GO enrichment analysis of the genes with high expression of ILC3a. G: Violin plots of the ILC3c highly expressed genes. H: Violin plots of the ILC3d highly expressed genes. I: Results of the GO enrichment analysis of the genes with high expression of ILC3d.

Furthermore, GO enrichment analysis of ILC3a cells showed that they mainly play a regulatory role in BP (biological process) enrichment (Figure S2E) for entries such as positive regulation of multicellular organismal processes, positive regulation of biological processes, positive regulation of developmental processes, and regulation of molecular functions. There was no significant difference between ILC3b cells and high-response genes. ILC3c cells overexpress the receptor gene FCGR3A encoding the Fc of immunoglobulin G, which has the function of regulating NK cells^52^; IL-17F is an effector cytokine of the innate and adaptive immune system and participates in antibacterial host defense and the maintenance of tissue integrity^53^. The protein encoded by the CD74 gene is related to class II major histocompatibility complex (MHC) and is an important chaperone for regulating antigen presentation of the immune response. ZBTB16 is an important transcriptional repressor gene involved in cell cycle processes and organism development (Figure 3F)^54^. ILC3d cells highly expressed the tumor suppressor genes SPINT2 and the involved cellular defense gene NCR2. Moreover, the important chemotactic receptor gene GPR183 was expressed in all ILC3 subsets (Figure 3G). The results of GO enrichment analysis of the ILC3d subset show that it mainly plays a role in regulating the biological process and development of cells in BP, with enrichment (Figure S2F) of terms such as singular organic process, regulation of multicellular organic development, and negative regulation of the biological process.

We have successfully described the typing and the localization of different cells in the pig gut from a transcriptomic landscape through the different expressions of genes among different cells. However, in terms of protein expression, we still need to perform certain experiments to confirm the porcine intestinal innate immunity typing and to detect protein targets. Thus, a monoclonal antibody against porcine RORC was generated and validated by flow cytometry. Then, we detected the subsets of ILCs by flow cytometry (Figure S4A), which was consistent with the expression of scRNA-seq. ILC3s were most abundant among all ILCs in the lamina propria of the porcine jejunum, accounting for approximately 80%, while the proportions of ILC2s, ILC1s, and NK cells were relatively low, and the expression of GATA3 in ILC3s was low. At the same time, through flow analysis of ILCs in the ileum, colon, lung, and blood, we found that ILCs existed in various tissues of pigs, and the composition of ILCs in different parts was quite different. In the small intestine, including the jejunum and ileum, ILC3s were dominant, while ILC1s/NK cells were dominant in the lung and blood (Figure S4B).

### 4. Pseudotemporal analysis of ILCs

ILC subsets develop from specific progenitor cells^55^; however, the fate of ILC subsets is not fixed. The environment regulates the transdifferentiation of ILCs through epigenetics^56^, contributes to the plasticity of ILC subgroups, and changes their functional polarization to adapt their response to different tissues and pathogenic stimuli^57^. Our results show that there is transdifferentiation among the intestinal ILC subgroups of pigs under certain conditions, and there are different enrichment positions in the whole pseudo-time-series trajectory, which suggests that there may be a transdifferentiation relationship among our ILC subgroups, among which most ILC3 subgroup cells are enriched in the early stage of the pseudo-time-series trajectory, especially ILC3b subgroup cells, which are enriched in the early stage of development. ILC1s, ILC2s, and NK cells were enriched at the late stage of the pseudotime trajectory (Figure 4A), which indicated that ILC3s in our gut had high plasticity and could be transformed into ILC1s, ILC2s, and NK cells. Because cells may differentiate into different types, there are many branches, and the differential genes between different branches may be the key to determining the fate of cell differentiation. Through the analysis of the branch nodes in the whole pseudotime sequence process, we found that there are three differentiation trends at the two differentiation nodes: the first trend is differentiation into ILC1 and NK cell subsets, the second trend is differentiation into the ILC2 subgroup, and the third trend is differentiation into the ILC3 subgroup. We express the differences in the differentiation of nodes in different directions in Figure 4B.

**Fig. 4.**
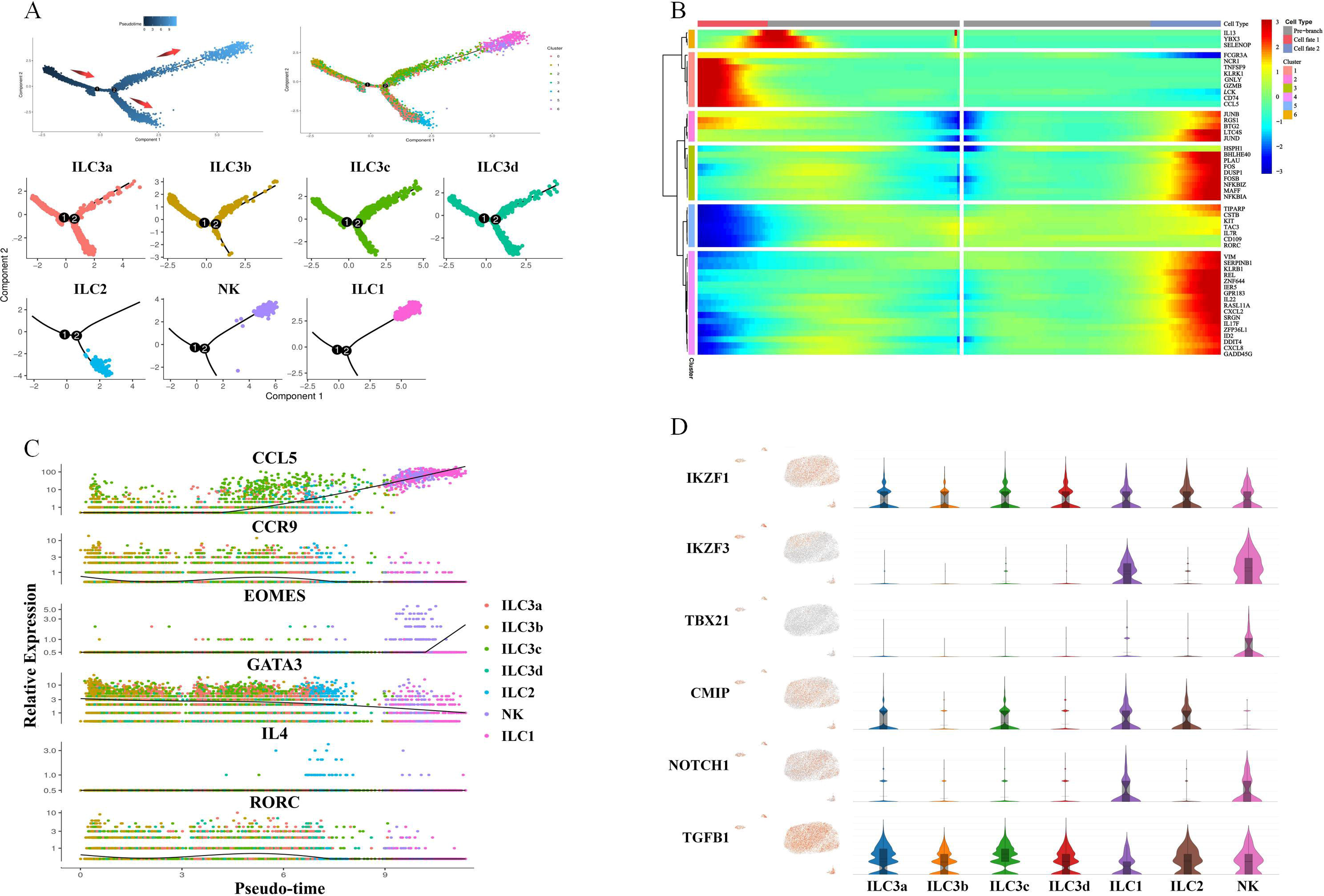
Pseudotemporal analysis of ILCs. A: Pseudotime traces created by Monocle 2 using the 7 clusters of ILCs. Arrows indicate the direction of differentiation, Show the enrichment position of each cluster in the Pseudotime(Different colors represent the different clusters), and black circles mark positions that are possible nodes of Pseudotime. B: Gene heatmap of cell differences between different branches. The top is the node branch in the pseudo temporal order, each row represents a gene, each column is the pseudotime point, the color represents the average expression value of the gene at the current time point, and the color is gradually reduced from red to blue, Genes with similar expression trends will cluster together to form a different cluster. C: Statistical map of the changes of marker genes in ILC clusters with the Pseudotime. D: For some genes that play regulatory transdifferentiation roles in subsets of ILCs, Violin plots of the genes regulating the transdifferentiation.

We found that the changes in some important landmark genes in each ILC subgroup during the pseudotime process, such as the highly expressed genes CCL5 and EOMES and other genes of NK cells and ILC1s, increased with the pseudotime response. The symbolic gene RORC of ILC3s changes with the pseudo-chronological process (Figure 4C). At the same time, we found some important expression levels (Figure 4D) that regulate the transdifferentiation genes of ILCs. It has been reported that Aiolos (IKZF3) (a TGF-β imprinted molecule) and TBX21 jointly promote the transdifferentiation of ILC3s to ILC1s^58-59^. We found that IKZF3 and T-bet are mainly expressed by ILC1 and NK cells in ILCs. We found that the responses of IKZF3 and IKZF1 promoted the differentiation of ILC3s into ILC1s and NK cells and promoted the transcriptional response of related genes^60^. Vonarbourg et al. found that IL-12 and IL-15 accelerated the conversion of ILC3s into ILC1s^61^. The C-Maf (CMIP) gene regulates ILC3s and can restrict the transdifferentiation of ILC3s to ILC1s^62^. In our results, some ILC3 subgroups (ILC3a and ILC3c), ILC1s, and ILC2s mainly expressed CMIP. The IL-4 gene secreted by ILC2s can also promote the transdifferentiation of ILC3s into ILC2s and maintain the identity of ILC2s^63^. T-bet and Notch signals from NK cells and ILC1s can drive NCR^−^ ILC3s to NCR^+^ ILC3s for ILC3 subgroup cells^64^. TGFβ is expressed in ILCs. Studies have shown that this gene can transform NCR^+^ ILC3s into NCR-ILC3s and can also induce ILC3s to differentiate into ILC regs and c-Kit^-^ ILC2s into c-Kit^+^ ILC2s^65^. It has also been found that RORγt-dependent ILC1s may be transdifferentiated from ILC3s in the presence of IL-12^66^. Although our results show that there is transdifferentiation of ILCs in the porcine intestine, much research data may be needed to prove the specific transdifferentiation direction among ILC subgroups in the porcine intestine and the related molecules that mediate this process.

### 5. Porcine and human ILC clusters are transcriptionally conserved but distinct from ILC subsets in mice

Through the analysis of ILCs in the human intestinal lamina propria and those in the mouse intestinal lamina propria, we found that ILCs in the human intestinal tract were mainly ILC3s, but a small number of ILC1s were found, whereas no ILC2s or NK cells were found. Certain proportions of ILC2s and ILC3s were found in mice, but ILC1s and NK cells (Figure S5A) were not found. Cluster analysis of ILC subsets in the intestinal tract shows that ILC3s in the human intestinal tract can be divided into four subgroups, namely, ILC3a, ILC3b, ILC3c, and ILC3d, and we show the genes with significant differential expression in each ILC3 subgroup and the ILC1 group (Figure S5B). ILCs in the porcine intestine are mainly ILC3s (Figure S5C). We found that the proportion of ILC3s in the intestinal tract of mice was still high, but there was also a high proportion of ILC2s (compared with humans and pigs). We showed the genes with significant differential expression in each ILC3 subgroup and the ILC2 group (Figure S5D).

To compare the transcriptional landscape of ILCs in the intestinal lamina propria among different species, we integrated our ILC dataset with published ILC scRNA-seq data from healthy mice^67-68^. We found that the similarity of gene expression in ILCs of human-pig-mouse was high, with 9017 common genes. There were 1425 unique coexpressed genes between humans and pigs, 478 between humans and mice, and only 281 between pigs and mice. At the same time, we found that there were 842, 524, and 290 genes expressed by humans, pigs, and mice, respectively (Figure 5A). Although we found a high similarity between human-pig-mouse, the abundance of gene expression of ILCs within the three species was quite different (Figure 5B) (Table S6).

**Fig. 5.**
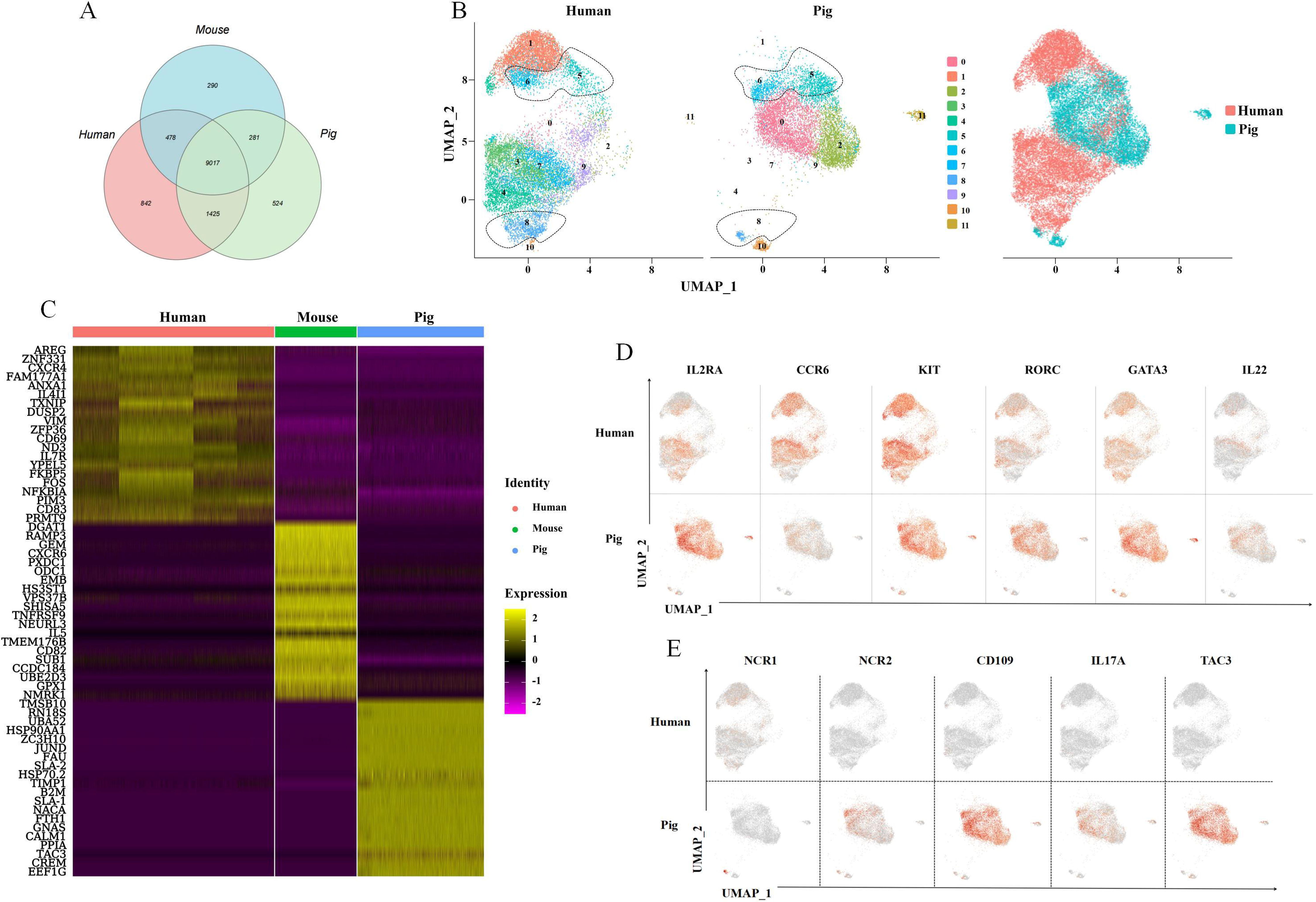
The Pig and Human clusters of intestinal ILCs are transcriptionally conserved. A: Venn statistics of genes expressed in human-pig-mouse intestine ILCs (Statistical mapping after normalization according to the number of expressed gene names). B: Statistical heatmap of the top 20 highly expressed genes in human, pig, and mouse intestine ILCs. C: UMAP plots showing the immune landscape of human and pig intestine ILCs enterocyte cluster subsets(And were divided into 12 clusters). UMAP with color coding according to donor origin. D: 6 marker genes coexpressed in human and pig ILCs cells are shown. E: 5 marker genes are expressed differently in human and pig ILCs cells.

By performing unbiased cluster analysis between human and pig ILCs, we found that clusters 8 and 10 were NK and ILC1 subgroups, cluster 11 was the ILC2 subgroup, and the others were ILC3 subgroups (clusters 0, 1, 2, 3, 4, 5, 6, 7, and 9) (Figure 5C). Some ILCs clustered with the pig intestinal tract, including the ILC3 subgroup, clusters 5 and 6, while human ILC1s clustered with pig NK cells. We found some similarities in the gene expression of ILCs between humans and pigs, such as the expression of the ILC signature genes IL-2RA, CCR6, KIT, RORC, and GATA3 (Figure 5D). At the same time, we found the differential expression of some genes, such as the expression of NCR1 in human ILC3s, which was greater than that of NCR2 in pig ILC3s (Figure 5E). CD109 had high expression in pigs but hardly any expression in humans. The expression level of the IL-22 gene is similar to that of human ILC3s, but the expression level of IL-17 in porcine ILC3s is higher than that of humans. The expression level of the TAC3 gene in porcine ILC3s was very high, but there was no expression in human ILC3s.

For cluster 5, cluster 6, and cluster 8, which cluster humans and pigs together, we found that humans and pigs highly coexpressed genes in cluster 5, including HSPA6, BAG3, DNAJB4, ZFAND2A, CKS2, and others (Figure S6A) and had a higher enrichment in BP and CC (Figure S6B). The highly expressed genes in cluster 6 included ZFP36L1, ST8SIA4, F2R, SLC8A1, and TGFBR2 (Figure S6C), and there was a higher enrichment in BP and CC (Figure S6D). The highly expressed genes in cluster 8 included SAMD3, NKG7, IRF8, SYTL3, and ADGRE5 (Figure SE), and there was a higher enrichment in CC and MF (Figure S6F).

The intestinal ILCs of mice were detected by flow cytometry. After CD45^+^LIN^-^CD127^+^ cells were gated, we detected the expression levels of RORγt and GATA3 and found that the ILC3:ILC2 ratio was approximately 2:1 (Figure 6A), which was consistent with the scRNA-seq data. The cluster analysis results showed that the gene expression of ILCs between mice and pigs was quite different compared to humans versus pigs, and subgroups were divided into 12 clusters, but almost no common cluster subgroup was found. There was an NK subgroup (cluster 12), an ILC1 subgroup (cluster 0), an ILC2 subgroup (clusters 7, 8, and 11), and an ILC3 subgroup (clusters 0, 1, 2, 3, 4, 5, 6, and 9) (Figure 6B). We found that the expression levels of the hallmark genes of ILCs between pigs and mice were similar, such as IL-7Rα, IL-23R, GATA3, RORC, and ID2. Among them, IL-23R and RORC were expressed by ILC3s, while ILC2s expressed higher levels of GATA3 (Figure 6C). At the same time, we found that the cytokines IL-4, IL-5, and IL-13 in the ILC2 subgroup were more strongly expressed in mouse ILC2s, while CXCL14 was more strongly expressed in porcine ILC2s (Figure 6D). The transcription of the cytokines CXCL2, CXCL8, IL-22, TIMP1, TNF, ZBTB16, and other genes in the ILC3 subgroup was more intensively expressed in porcine ILC3s (Figure 6E). For example, CD90 (THY1), a member of the superfamily encoding cell surface glycoprotein and immunoglobulin, is only expressed in mouse ILCs. The expression level of AHR is the highest in mouse intestinal ILC2s. The mouse ILC3s highly expressed the CX3CL1, cytokine signaling inhibitor gene SOCS3, and chemokine receptor gene CCR7 (Figure 6F).

**Fig. 6.**
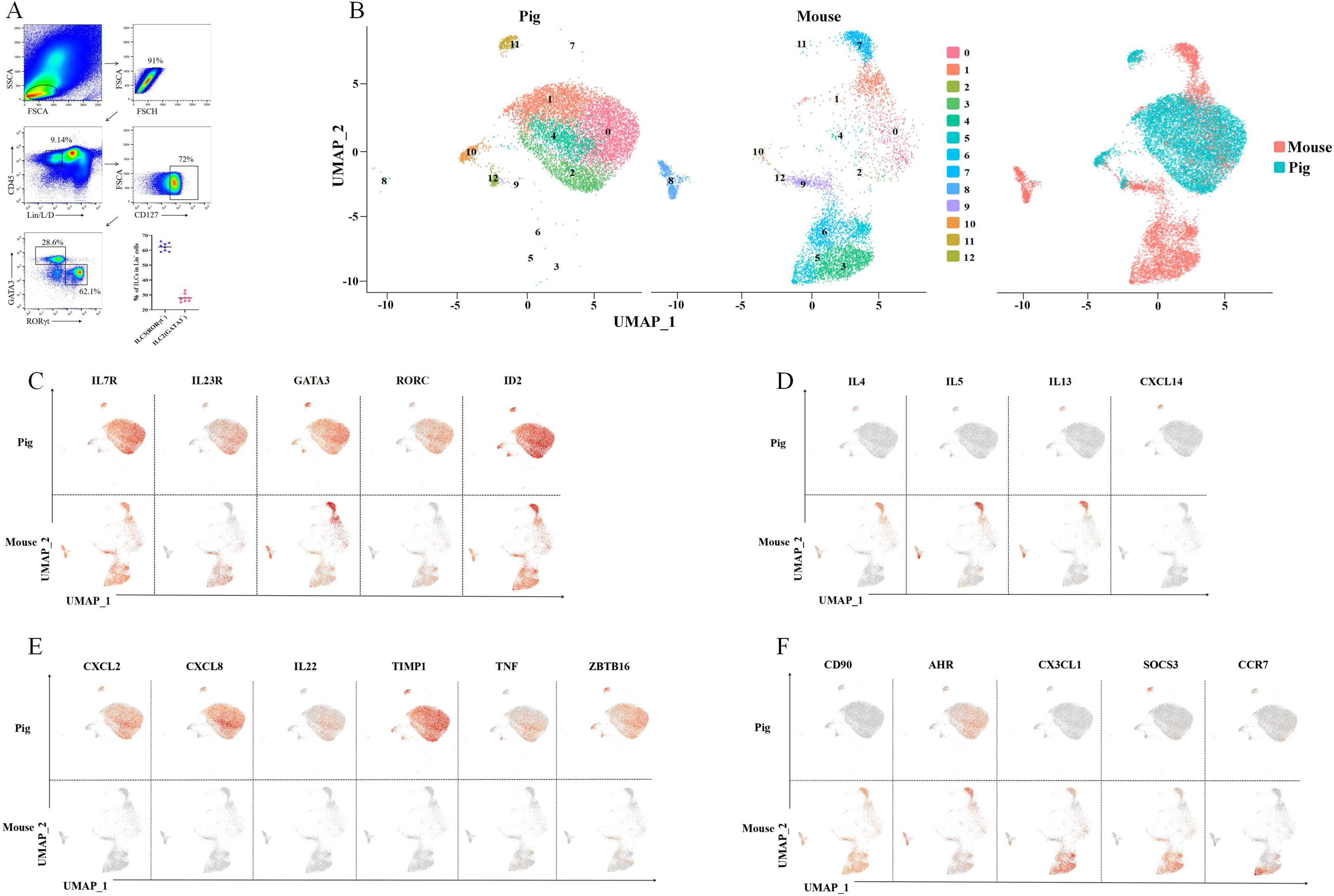
The Pig and Moues clusters of intestinal ILCs express differential genes. A: Mouse intestinal ILCs were examined by flow cytometry. B: UMAP plots showing the immune landscape of human and mouse intestine ILC clusters. C: 6 marker genes coexpressed in pig and mouse ILCs cells are shown. D: 4 cytokine marker genes’ differential expression in pig and mouse ILC2 cells are shown. E: Six cytokine marker genes are expressed differently in pig and mouse ILC3 cells. F: 5 marker genes are expressed differently in pig and mouse ILCs cells.

According to our human-pig-mouse coclustering analysis, the gene expression species similarity of intestinal ILCs in humans, pigs, and mice was high. However, the gene expression abundance of ILCs varies greatly between species. For the clustering of ILCs between humans and pigs, we found common-clustered ILC3 subsets and human ILC1s were also clustered together with porcine NK cells. However, the intestinal ILCs between pigs and mice differed and did not cluster obviously.

## Discussion

We sorted 200,000 L/D^-^LIN^-^CD45^+^ cells from porcine jejunum LPLs by flow cytometry for scRNA-seq and finally obtained 15,763 translational landscapes after filtering. We found helper-like ILCs (including ILC1s, ILC2s, and ILC3s). Interestingly, we performed a more detailed analysis of ILCs in jejunal LPLs. We found that porcine intestinal ILCs mainly included four groups, namely, ILC3s, ILC1s, ILC2s, and NK cells (sorted according to the proportion of cell number). Among them, ILC3s were the main ILC in the pig jejunal LPLs, and a further subset analysis of ILC3s based on the gene expression profiles revealed four subsets of ILC3s and identified transdifferentiation among subpopulations of ILCs. However, we did not find LTi cells in the intestinal ILCs of pigs, there were no clustered subgroups, and no gene list of LTA (the marker gene of human LTi cells) was found.

In the porcine jejunum, CD2, as a broad-spectrum and relatively stable whole T-cell antigen, is expressed not only in mature T cells and thymocytes but also in ILCs^69^. The jejunal ILCs are mainly ILC3s and have signature transcribed genes similar to human genes such as IL-23R, hydrocarbon receptor (AHR), IL-7Rα, NCR2, RORC and GATA3 (Figure 2E, G). Expression of the highly expressed aryl AHR of ILC3s can regulate the ILC balance and control chromatin accessibility at the AHR locus through positive feedback^70^. The host can also regulate the secretion of IL-22 by ILC3s by participating in the AHR pathway, thereby generating appropriate immunity against various pathogens^71^. AHR expression is also important for maintaining NKp46^+^ ILC3s^72^. Studies have found that ILCs expressing NKp44 mainly develop into ILC3s^73^. IL-7-dependent maintenance of ILC3s is required for the normal entry of lymphocytes into lymph nodes^74^. Runx3 is required for the normal development of both ILC1s and ILC3s and controls the survival of ILC1s. Runx3 is also required to express the transcription factor RORγt and its downstream target transcription factor AHR in ILC3s^75^. At the same time, we found that ILC3s have rich subtypes, such as ILC3a with high expression of CXCL2 and IL-22, ILC3c with high expression of IL-17F, and ILC3d with high expression of CD74 (Figure 3C). We also found that ILC3s have the potential to differentiate into ILC2s, ILC1s, and NK cells. Although ILC3b cells have no obvious highly expressed genes, they have more obvious plasticity in the transdifferentiation among ILC subgroups. At the same time, we found some genes related to this process, such as IKZF1 and TGFB1, that were coexpressed by ILCs. ILC1s and NK cells express IKZF3, TBX21, and Notch, and ILC3s, ILC2s, and ILC1s express CMIP. These genes play an important role in regulating the transdifferentiation of ILCs.

The difference between NK cells and ILC1s is usually the expression of the transcription factor exoderm protein (Eomes)^76^. ILC1s and NK cells have overlapping phenotypes and functions but depend on different developmental pathways. NKG7, CCL5, and LITAF showed a higher expression in both ILC1 and NK (Figure 2F). The transcriptional repressor of ILC1s, MXI1, can negatively regulate MYC function and is a potential tumor suppressor (Figure 2G). For the marker genes of NK, such as LYST, ISG15, and GIMAP4T (Figure 2H), some studies have reported that LYST regulates lysis granules in cytotoxic T cells and NK cells^77^. ISG15 is a ubiquitin-like protein gene important in innate immunity^78^. GIMAP4T plays an important role in regulating the survival and development of lymphocytes^79^. The hallmark receptor gene PTGDR2 for ILC2s can mediate the proinflammatory chemotaxis of eosinophils, basophils, and lymphocytes produced during allergic inflammation^80-81^. IL-4, IL-5, IL-13, and CSF2, with high expression in ILC2s (Figure 2F), belong to the same gene cluster and are located on the chromosome 5q31 region, which is related to the differentiation of Th2 cells/ILC2s^82^. Among the transcription factors of ILC2s, GATA3 is an important regulator of the cell development of ILCs and is also very important for developing ILC3s^83^. RORA has been proven to contribute to the transcriptional regulation of some genes involved in circadian rhythm and is important for developing ILC2s and ILC3s and regulating macrophages and Treg cells^84^. ZBTB16 encodes a zinc finger transcription factor and participates in the progression of the cell cycle^85^. The single-cell map of lung ILC2s shows that the tissue-resident ILC progenitor cells (ILCPs) and effector cells heterogeneously express ZBTB16. The YBX3 gene negatively regulates the cell response to tumor necrosis factor and programmed cell death^86^. In steady-state tissues, ILC2s are more recognized as a major source of type 2 cytokines than Th2 cells and are a key driver of allergic reactions in the early stages of the disease^87^.

Compared with pig and mouse ILCs, human and pig intestinal ILCs are transcriptionally conserved. ILCs were mainly ILC3-predominant in humans and porcine intestines, with an almost complete absence of ILC2s, while ILC2s were present in high proportions in mice. We also found some ILC marker genes commonly expressed in humans, pigs, and mice. The transcription factor GATA3 is essential for developing all ILCs, and studies have reported that GATA3 can not only control IL-7Rα but can also regulate the transcription factors TBX21 and RORC^88^. Some transcription factors, such as RORC, TBX21, and Notch (Figure 2G), are essential for differentiating ILC subgroups. In the differentiation of human ILCs, some studies have found that the Notch→RORC→IL-23R pathway plays an important role, and these genes are expressed similarly in pig and human intestinal ILCs^89^. At the same time, there is some specific expression of genes in porcine jejunum ILCs (Figure 5E). For example, in porcine ILCs, the NCR1 gene is only expressed in NK cells but not in other ILC subgroups, and porcine ILC3s showed a higher transcription level of the IL-17 gene. We also found the SLA1 gene, which specifically responds to porcine ILCs and regulates the signal transduction of the T-cell receptor (TCR).

In summary, the results of cross-species analysis and comparison of ILCs have laid a foundation for further exploration of pigs as a human medical model^90^. We found a high similarity of intestinal ILCs between pigs and humans, confirming the feasibility of pigs as a human surrogate model for studies related to intestinal mucosal immunity. Especially in research concerning the cell target of ILC-related and intestinal mucosal immune diseases such as enteritis and inflammatory bowel disease (IBD), pigs have a natural advantage as model animals.

Studies have reported the existence of regulatory relationships such as the gut-brain axis^91^, gut-lung axis^92^, and gut-kidney axis^93^. In our results, porcine ILC3s were found to express the TAC3 gene. The TAC3 gene encodes neurokinin B (NKB), whose receptor is NK3R^94^, and the combination of NKB and NK3R can promote gonadotrophin-releasing hormone (GnRH) release^95^. GnRH binds to specific receptors on the pituitary gonadotropin to promote the release of sex hormones. Therefore, TAC3 plays an important role in the gonadotrope axis, and we proposed the gut-gonadal axis, where porcine intestinal ILC3s regulate gonadal development.

Our research on ILCs in porcine intestines can promote the intestinal health of pigs, and at the same time, it is of great significance to biomedical research, animal models, and global food health. However, there is little biological information due to the limited studies on the intestinal tract of pigs, especially ILCs, a new lymphocyte family. Our discovery is the first single-cell description of the ILC transcriptome landscape in the pig jejunum, focusing on research on ILCs in the jejunum. In addition, other lymphocyte types in the intestine also play an important role in intestinal health, but they are outside the scope of this sudy. Our work is based on the intestinal health of the jejunum, identifying the transcription information of ILCs in the jejunum by scRNA-seq and then inferring the functional subgroup. This work also provides more biological knowledge related to this species of pig.

Although our data can be used as favorable evidence for the existence of ILCs in porcine intestines and can help us to explore which genes the ILCs express to play an immunomodulatory role and how they transdifferentiate under complex environmental conditions in the intestines, this understanding is only applicable to the intestines at present, while the roles of ILCs in other tissues of pigs are still unclear. Therefore, our data provide an exciting starting point for the study of porcine ILCs, and it is necessary to further analyze ILCs in other parts of pigs in the future to analyze the tissue heterogeneity and developmental characteristics of porcine ILCs.

## Limitations of the study

Although this study provided a comprehensive transcription analysis of ILCs in the lamina propria of the porcine intestine, we only analyzed single-age/sex pigs. In this study, to obtain more ILCs, we enriched lymphocytes in the lamina propria with Lin^-^, which led to the loss of many T/B lymphocyte groups and nonlymphocytic groups. For example, the CD8 and CD4 antibodies we used could lead to the loss of NK and LTi cells. In addition, although we found that there is a great transcription difference between the porcine intestinal ILC subgroup and mouse intestinal ILC subgroup, there is a high similarity with human intestinal ILCs, and we cannot rule out that the differences among human, porcine, and mouse species are due to biological and technical effects, such as differences in physiological age, tissue preparation methods, and sequencing saturation. In the future, if more reagents or better methods are available, we should use various methods to verify our findings.

## Materials and Methods

### Animals and sample collection

All Specific pathogen Free (SPF) experimental pigs were purchased from Harbin Veterinary Research Institute, and all piglets were weaned at 3 weeks old. When piglets were 4 weeks old, we used flow cytometry to analyze the immune cells of the jejunum lamina propria. Then, we collected samples from the jejunum of 4 SPF female pigs, collected cell samples by flow sorting, performed scRNA-seq, and made sections of tissue samples for immunofluorescence experiments. All animal experiments met the requirements of the Animal Management and Ethics Committee of Jilin Agricultural University.

### Cell separation

For cell isolation of the jejunum, immediately after the euthanasia of piglets, approximately 5 cm of the jejunum was collected and stored in phosphate-buffered saline (PBS) at 4°C. The outer muscle layer was peeled off, and the intestine was cut longitudinally to expose the lumen. The tissue was rinsed gently with PBS to remove the intestinal contents and cut into 1 cm large and small intestine segments with scissors.

The tissue was transferred to the separation solution (15 mL of RPMI-1640, 1% double antibody, 1% HEPES, 5 mM EDTA, 2 mM DTT, and 2% heat-inactivated fetal bovine serum (FCS)) and incubated in a shaking incubator at 37°C and 200 rpm for 28 minutes. After shaking for 2 minutes, the remaining intestinal tissue was retained, the washing liquid was removed, and PBS at 4°C was used for washing.

The remaining tissues were transferred to the digestive juice solution (8 mL RPMI-1640 medium, 1% double antibody, 1% HEPES, 50 mg collagenase IV, 1 mg DNase I, and 2% FCS) and incubated in a shaking incubator at 37°C and 250 rpm for 20 minutes. Digestion was continued for 20 minutes, after which the samples were shaken vigorously for 2 minutes, and then the digestive juice was collected through a 70-µm cell membrane.

Culture solution (10 ml RPMI-1640, 5% FCS) was added to the collected digestive juice, centrifuged at 4°C and 450xg for 8 minutes, resuspended in 40% Percoll solution, then slowly added to 80% Percoll solution, and centrifuged at 24°C and 500xg.

### Antibodies staining

Monoclonal antibody against porcine RORC for flow cytometry was produced by Jieyu Company (RORC Gene ID: 100622477) (clone 11F4). Hybridoma cells were preserved by the China Center for Type Culture Collection (CCTCC) (No. C2022219), and related products have been patented (No. 202210980598. X).

Antibodies for pigs were as follows: Invitrogen: CD2 monoclonal antibody (14-0029-82), CD3e monoclonal antibody (MA5-28774), CD3e monoclonal antibody (Biotin) (MA5-28771), CD11b monoclonal antibody (MA5-16604), CD11c monoclonal antibody (APC) (17-0116-42), CD21 monoclonal antibody (A1-19243), CD21 monoclonal antibody (PE) (MA1-19754), CD45 monoclonal antibody (FITC) (MA5-28383), CD117 (c-Kit) monoclonal antibody (APC) (17-1171-82), CD163 monoclonal antibody (PE) (MA5-16476), CD172a monoclonal antibody (MA5-28299), CD335 (NKp46/NCR1) monoclonal antibody (PE) (MA5-28352), SLA Class II DR monoclonal antibody (MA5-28503), GATA3 monoclonal antibody (PE-Cyanine7) (25-9966-42), T-bet monoclonal antibody (PE) (12-5825-82), and goat anti-mouse IgG H&L (PE-Cyanine5) (M35018).

BD Pharmingen: Fixable Viability Stain 780 (565388), purified rat anti-mouse CD16/CD32 (Mouse BD Fc Block) (553142), mouse anti-pig CD3ε (PE) (561485), mouse anti-pig CD4a (PE) (559586), mouse anti-pig CD8a (FITC) (551543), mouse anti-pig CD8a (Alexa Fluor 647) (561475), rat anti-pig γδ T Lymphocytes (PE) (551543), rat anti-pig γδ T lymphocytes (561486), and mouse anti-GATA3 (APC) (558686).

Abcam: goat anti-mouse IgG H&L (FITC) (ab6785), goat F(ab’)2 anti-mouse IgG H&L (PE) (ab7002), goat F(ab’)2 anti-rat IgG H&L (PE-Cyanine5) (ab130803), and rat anti-mouse IgG1 H&L (PE) (ab99605).

BioLegend: CD11b monoclonal antibody (APC) (101212), CD11c monoclonal antibody (PE) (301606).

R&D: porcine CD34 antibody (AF3890).

Antibodies for mouse were as follows: BD: Fixable Viability Stain 780 (565388), CD11b (Biotin) (557395), γδ T (Biotin) (553176), CD19 (Biotin) (553784), TCRβ (Biotin) (553168),

Ly6G/C (Biotin) (553124), TER-119 (Biotin) (553672), streptavidin protein (APC-cy7) (554063), CD45 (FITC) (551874), CD127 (PE-cy7) (560733), RORγt (PE) (562607), and GATA3 (Alexa Fluor 647) (560068).

### scRNA-seq

Enriched ILCs were obtained by flow sorting. We stained the total LPL extracted from the lamina propria with antibodies against LIFE, CD45, lin1 (CD3, CD21, γδT, CD11c, CD172a), and lin2 (CD4, CD8, CD163, CD11b) and then selected LIFE-CD45+Lin1-Lin2-cells by flow sorting. The concentration was 900 cells/μL, the cell viability was > 75, the total number of cells was adjusted to approximately 5×10^6^, and then SCRNA-SEQ was performed.

The cell suspension was loaded into Chromium microfluidic chips with 30 chemistry and barcoded with a 10× Chromium Controller (10× Genomics). RNA from the barcoded cells was reverse-transcribed, and sequencing libraries were constructed with Chromium Single Cell 30 reagent kit (10× Genomics) according to the manufacturer’s instructions. Sequencing was performed with Illumina (Novaseq PE150) according to the manufacturer’s instructions (Illumina).

### Data analysis

We used FastQC to perform basic statistics on the quality of the raw reads. Generally, celltanger count supports FASTQ files from raw base call (BCL) files generated by Illumina sequencers as input files. 10× Genomics® does not recommend additional processing of the sequence.

Raw reads were demultiplexed and mapped to the reference genome by the 10× Genomics Cell Ranger pipeline (https://support.10xgenomics.com/single-cell-geneexpression/software/pipelines/latest/what-is-cel l-ranger) using default parameters. Unless mentioned specifically, all downstream single-cell analyses were performed using Cell Ranger and Seurat. In brief, unique molecule identifiers were counted for each gene and cell barcode (filtered by CellRanger) to construct digital expression matrices. For secondary filtration by Seurat, a gene with expression in more than 3 cells was considered expressed, and each cell was required to have at least 200 expressed genes. Then, some of the foreign cells were filtered out.

### Secondary analysis of gene expression

Cellranger reanalyze takes feature-barcode matrices produced by cellranger count or cellranger agar and reruns the dimensionality reduction, clustering, and gene expression algorithms using cellranger default parameter settings.

The Seurat package was applied to perform data normalization, dimensionality reduction, clustering, and differential expression. We used the Seurat alignment method canonical correlation analysis (CCA) to analyze datasets. For clustering, highly variable genes were selected, and the principal components based on those genes were used to build a graph, which was segmented with a resolution of 0.6^25^.

To perform global analysis between samples, based on the filtered gene expression matrix by Seurat, between-samples differential expression analysis was carried out using the edgeR package to obtain zone-specific marker genes^96^.

For enrichment analysis of marker genes, Reactome pathway-based analysis of marker genes was implemented by the ReactomePA R package. REACTOME is an open-source, manually curated, peer-reviewed pathway database (https://reactome.org/).

The GSVA analysis was performed using the msigdb database (https://www.gsea-msigdb.org/gsea/msigdb/index.jsp)c2.cp.kegg.v7.4.symbols.gmt.

The results from CellDB were presented for the cell-type interaction network using iTALK software.

For trajectory analysis, Monocle2 was used to calculate the correlation of cells to obtain the minimum spanning tree, find the minimum path, project all other data points to the minimum path, and finally obtain the cell differentiation trajectory diagram^97^.

### Dataset integration

The porcine jejunal scRNA-seq data was derived from GSE175411^98^. We compared our data with published human intestinal ILC (GSA-Human: HRA000919) and mouse intestinal ILC (GSE166266) datasets. The Ensembl genome browser (Ensembl Genes 105) was used to convert human (GRCh38) and mouse (GRCm39) gene names to the corresponding pig gene names prior to integration (https://www.ensembl.org/biomart/martview/). The indiscriminate clustering algorithm was analyzed later^20^.

## DATA AVAILABILITY STATEMENT

The raw data for this article were deposited in the National Center for Biotechnology Information (NCBI) Sequence Read Archive (SRA) database under BioProject number PRJNA907920.

## AUTHOR CONTRIBUTIONS

Cell isolation, J.H.W., and M.Y.C.; data analysis, M.G, Y.S., and Y.Y.L; manuscript preparation and writing, J.H.W., Y.Z., X.X.L., and C.W.S.; supervision and project administration, C.F.W. and X.C. All authors contributed to the article and approved the submitted version.

## ACKNOWLEDGEMENTS

This work was supported by the National Natural Science Foundation of China (32273043, 32202890, U21A20261), the Science and Technology Development Program of Changchun City (21 ZY 42), the Science and Technology Development Program of Jilin Province. (20200402041 NC), and China Agriculture Research System of MOF and MARA (CARS-35). We thank Novogene Co., LTD for providing technical services.

## ETHICS

The animal management procedures and all laboratory procedures abided by the regulations of the Animal Care and Ethics Committees of Jilin Agriculture University, China.

## Conflict of Interest

The authors declare that the research was conducted without any commercial or financial relationships that could be construed as a potential conflict of interest.

This paper does not report the original code.

Any additional information required to reanalyze the data reported in this paper is available from the lead contact upon request.

## Figure legends

**Figure. S1.**
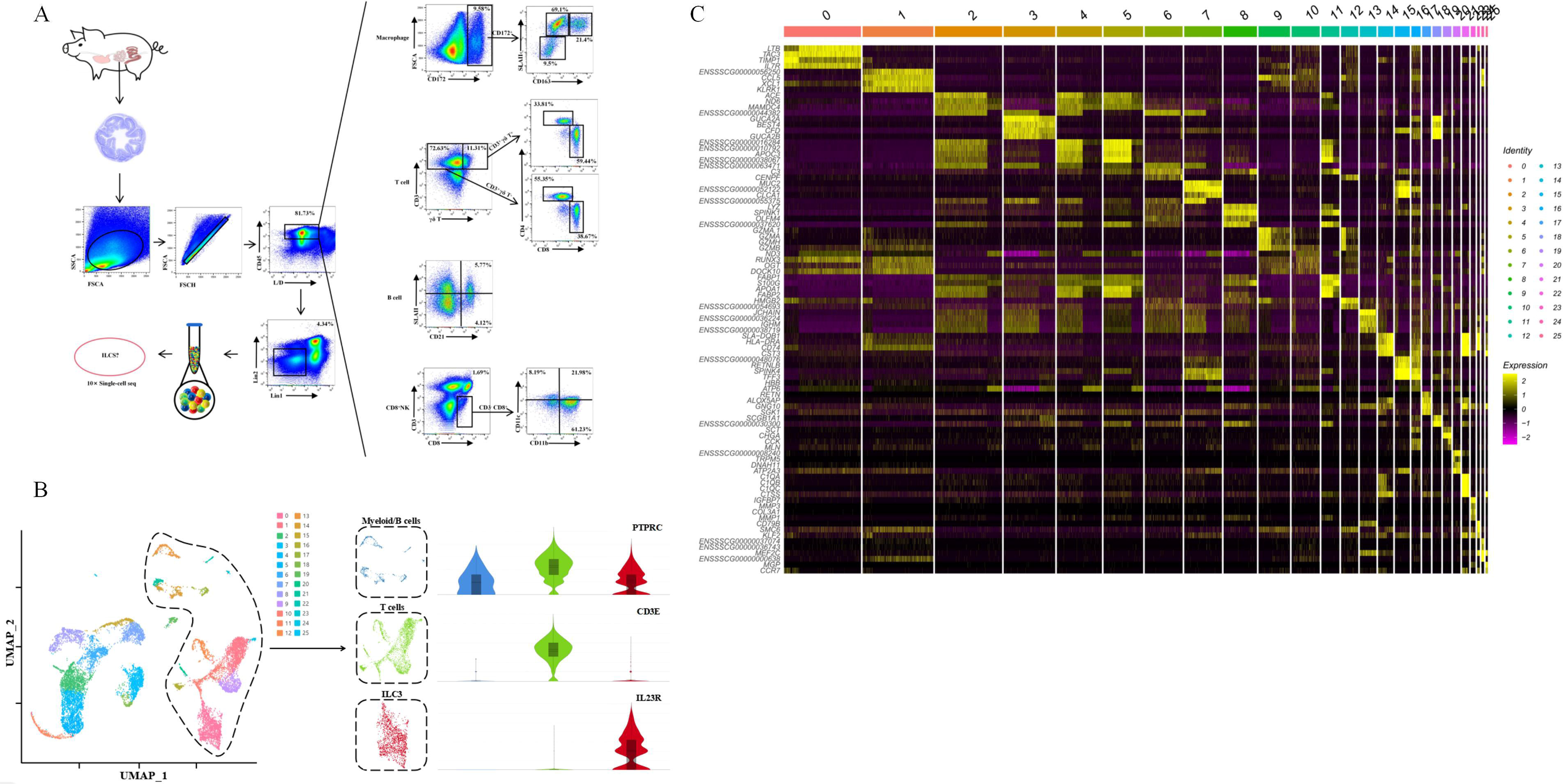
Identification of all cells in pig jejunum by flow cytometry and scRNA-seq. A: Flow cytometry analysis of immune cells in the lamina propria of the pig jejuna. Isolation of immune cells from the jejunal lamina propria shows cellular subpopulations by the signature cellular marker proteins. The ILCs can also be obtained by flow sorting for further single-cell analysis. B: Uniform manifold approximation and projection (UMAP) visualization of intestine cell types, colored by cell clusters. Clusters were identified using the graph-based Louvain algorithm at a resolution of 0.5. C: Heatmap showing row-scaled mean expression of each cluster’s four highest differentially expressed genes. Abscissa shows different clusters, and the ordinate shows differentially expressed gene names.

**Figure. S2.**
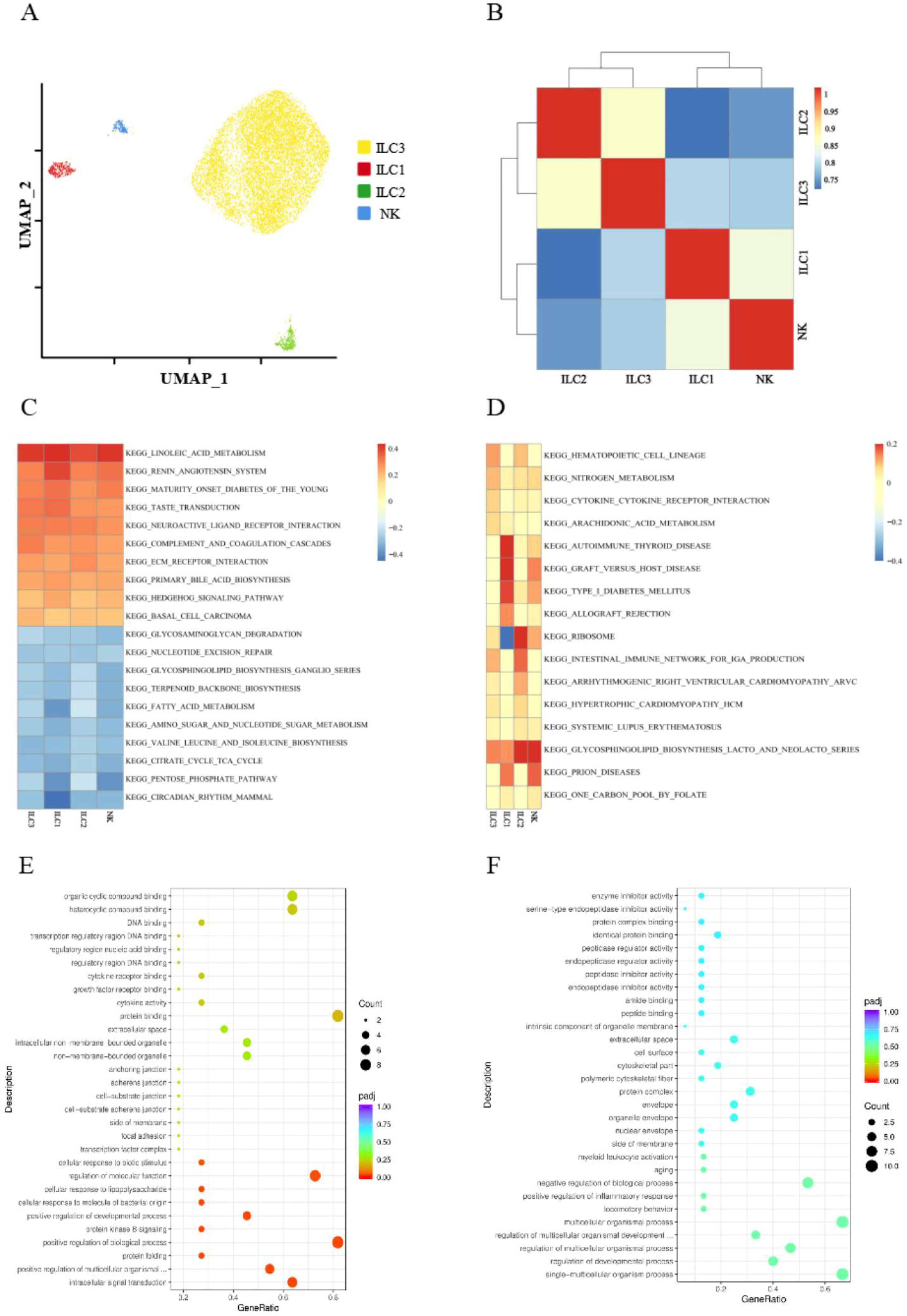
Further analysis of the four clusters of ILCs. A: UMAP plots showing the immune landscape of the ILC subpopulations (92.57% of ILC3, 2.99% of ILC1, 2.73% of ILC2, and 1.71% of NK). B: Gene heatmap of cell differences between branches, Genes with similar expression trends will cluster together to form a different cluster. Colors represent the degree of correlation between the ILC clusters. C: The GSVA heatmap is according to the default pathway order. The top 10 plots were selected, and the different colors represent the enrichment of each subset gene set in the pathway. D: GSVA analysis of the four subset gene sets of ILCs, heatmap showing the top four significantly enriched pathways.

**Figure. S3.**
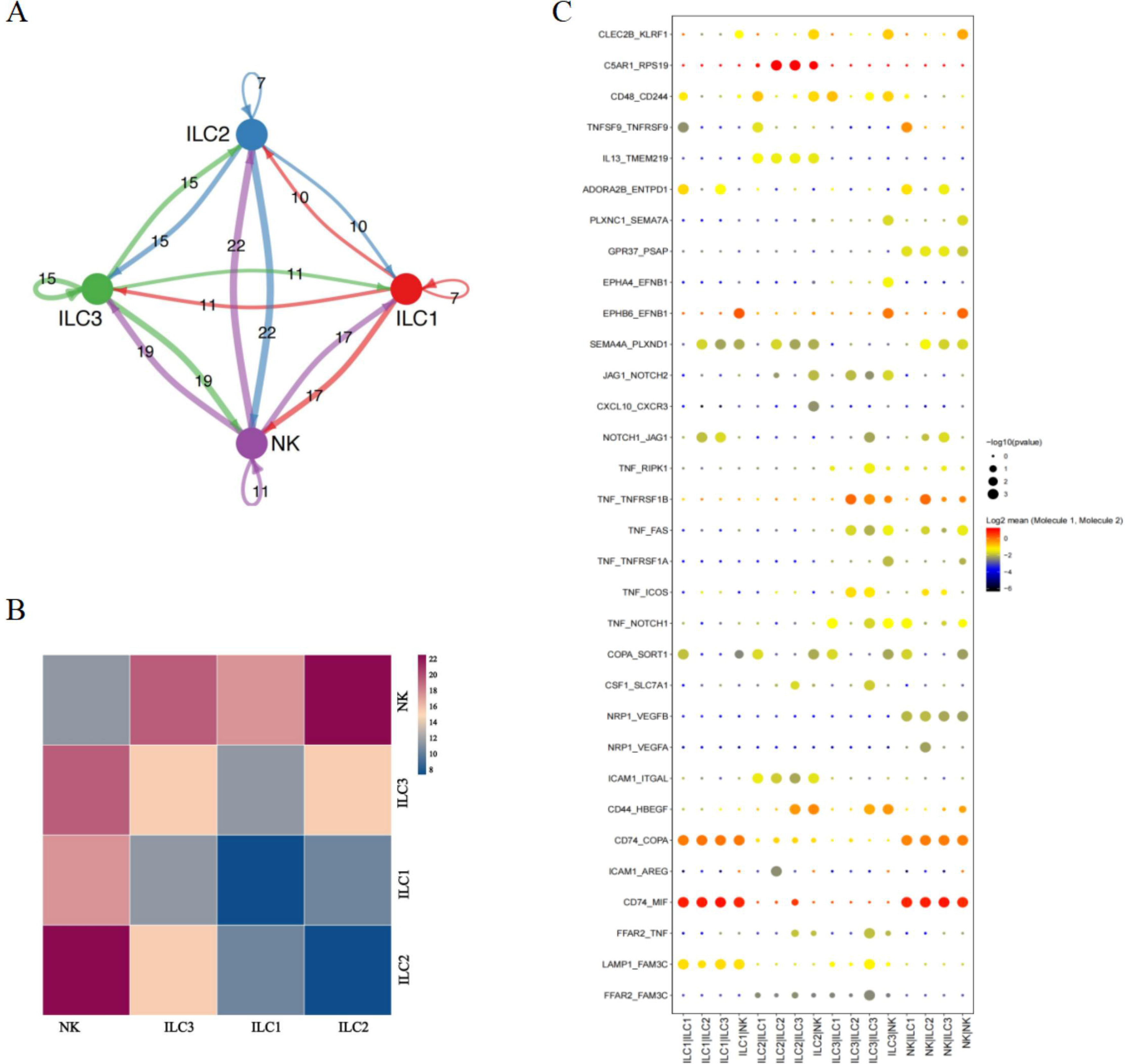
CellPhoneDB analysis of the four clusters of ILCs. A: Network of interacting relationships between cell types: where the nodes surface the cell type, in which the arrow represents the pointing relationship, and the number on the side represents the ligand-receptor logarithm (the more the number, the thicker the line) B: Heatmap of the number of ligands among cells in the ILC subsets. Figure. C: Statistical dot plot of significant target ligand genes among ILC subsets. The abscissa is the group of cells undergoing ligand interaction, and the ordinate represents the ligand information.

**Figure. S4.**
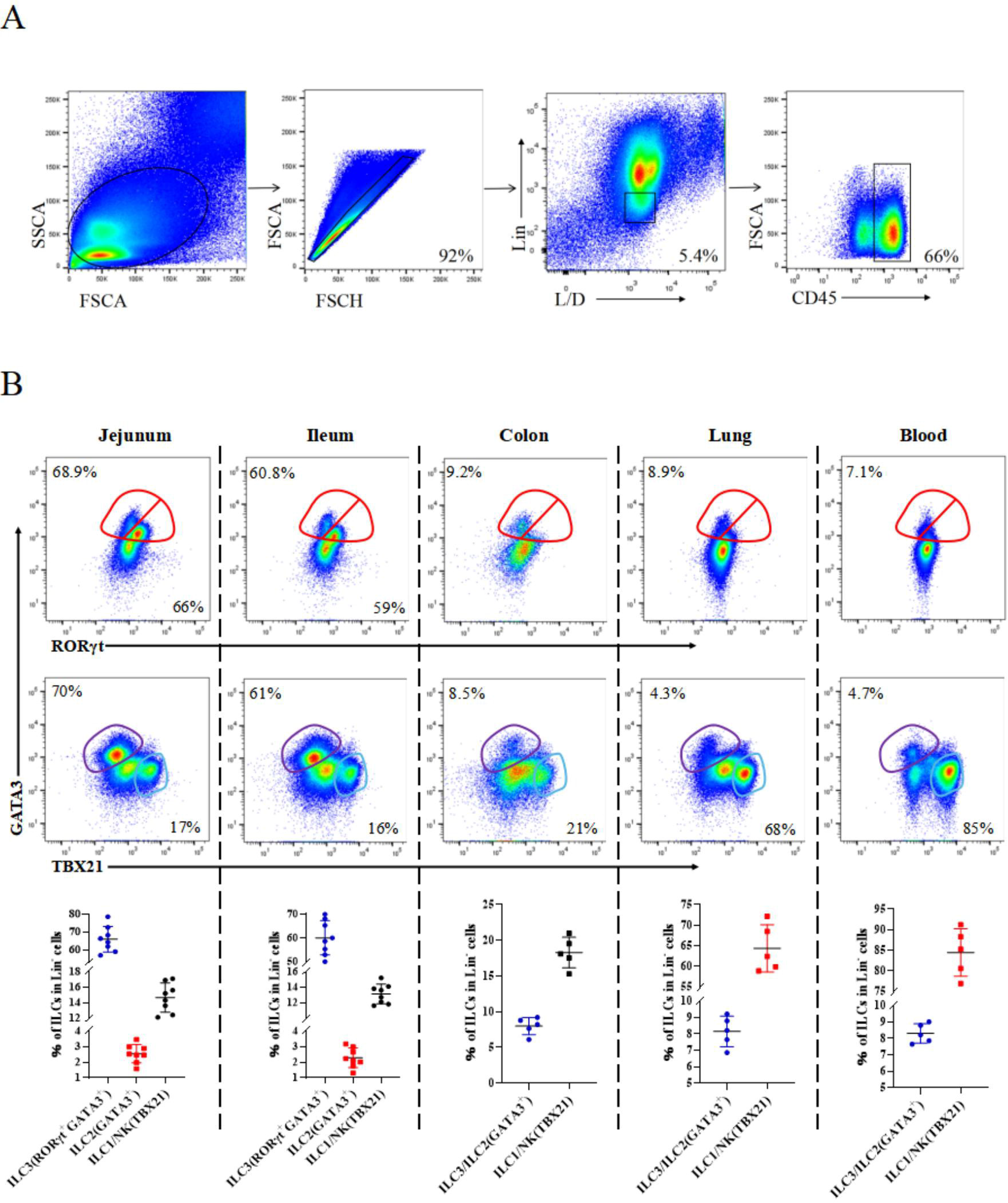
Flow cytometry analysis of intestinal ILCs, different tissues represent the tissue heterogeneity of ILCs. A: Schematic representation of the flow cytometry detection ring gate of cells in ILC subsets. B: Flow cytometry detection result statistics for ILC in different tissues, the proportion of ILCs in different tissues was arranged and displayed.

**Figure. S5.**
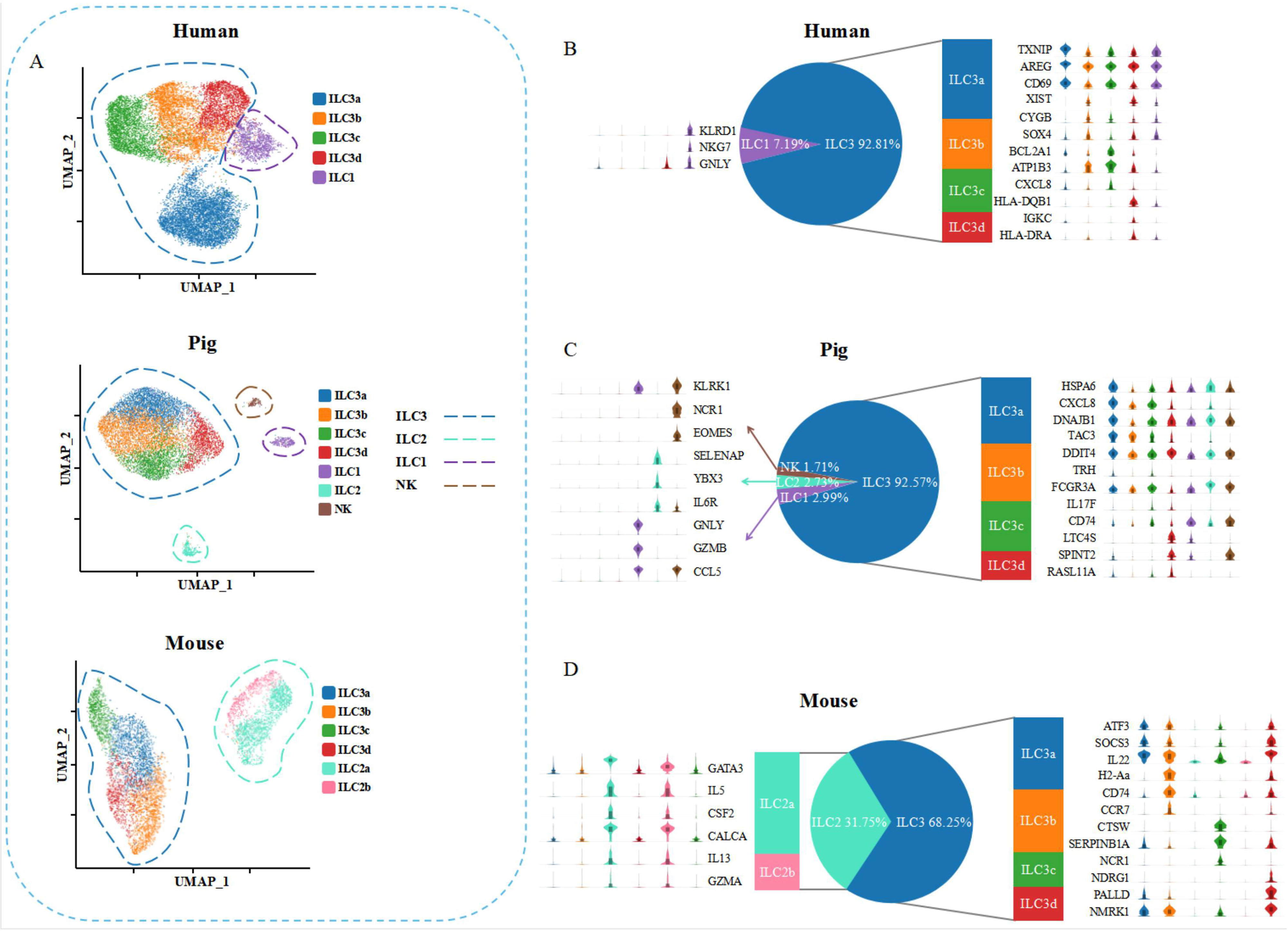
UMAP clusters analysis of intestinal ILCs in human, pig, and mouse, Statistical presentation of highly expressed genes in ILCs. A: UMAP plots showing the immune landscape of human, pig, and mouse intestine ILCs clusters. Dashed lines with different colors represent the positions of ILC3, ILC1, CIL2, and NK cells, respectively. B: Statistical analysis of human intestinal ILCs, separately displaying three highly expressed genes for each cluster. C: Statistical analysis of mouse intestinal ILCs, separately displaying highly expressed genes for each cluster.

**Figure. S6.**
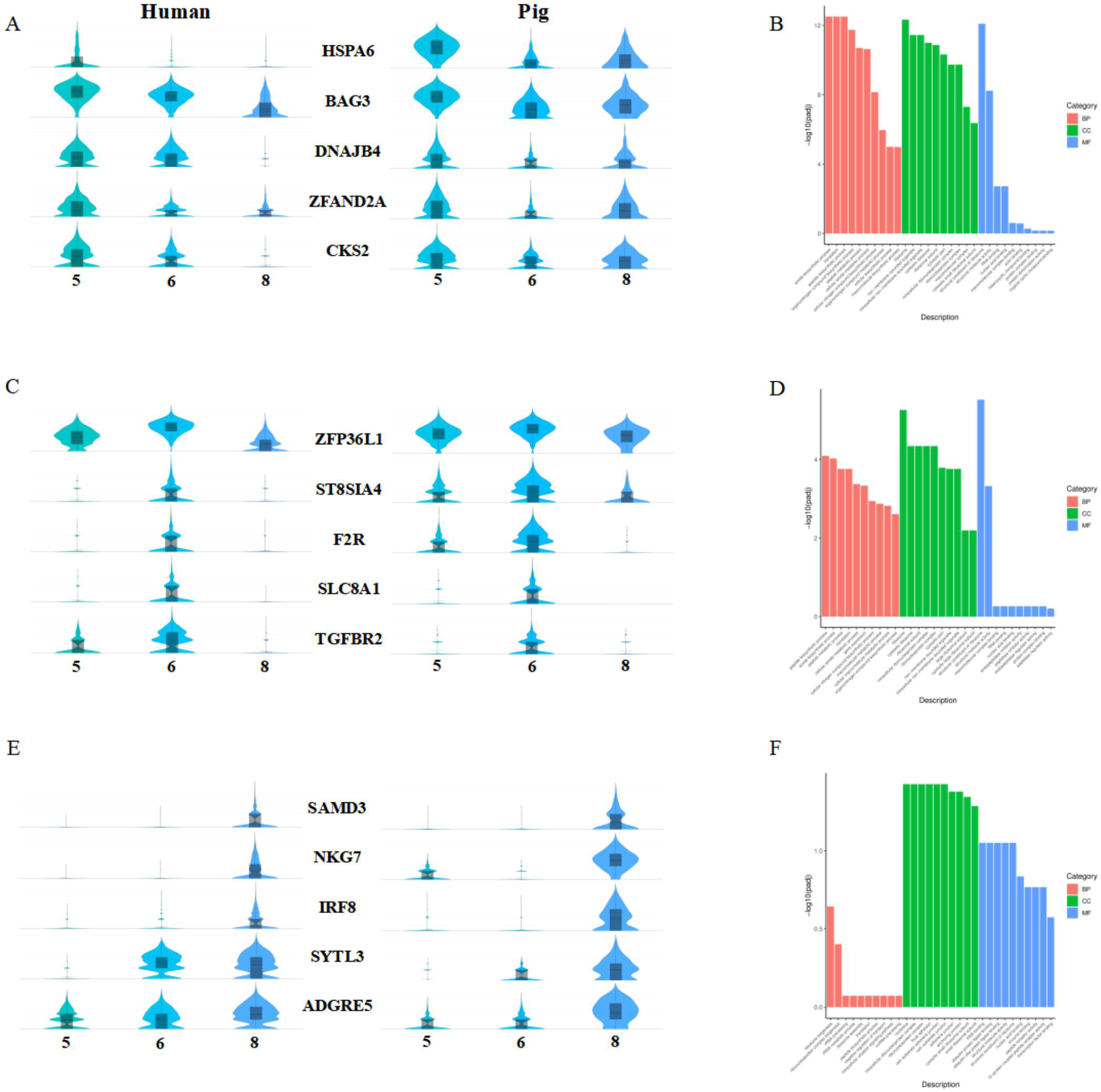
Three subpopulations of human and pig co-clustered were further analyzed. A: Cluster 5 Violin plots of human and pig significant table answer genes, showing the top five. B: Bar graph of GO enriched pathways for genes in Cluster 5. C: Cluster 6 Violin plots of human and pig significant table answer genes, showing the top five. D: Bar graph of GO enriched pathways for genes in Cluster 6. E: Cluster 8 Violin plots of human and pig significant table answer genes, showing the top five. F: Bar graph of GO enriched pathways for genes in Cluster 8.

